# Non-invasive Neuromodulation Targeting Approach by Mapping Stimulations and Lesions That Modify Visual Memory

**DOI:** 10.64898/2026.04.10.717784

**Authors:** Simon Kwon, Soyoung Lee, Joshua S. Siegel, Nicole Chiulli, Michael Freedberg, Melissa Hebscher, Joshua Hendrikse, Molly S. Hermiller, Gong-Jun Ji, Arielle Tambini, Eyre Ye, Shira Cohen-Zimerman, Maurizio Corbetta, Jordan Grafman, Joel L. Voss, Shan H. Siddiqi

## Abstract

Therapeutic brain stimulation is believed to target specific networks, but targeting approaches for memory remain debated. For other symptoms, neuromodulation targets have been localized by mapping connectivity of lesions and stimulation sites to specific symptoms. This approach has yielded networks for global memory, but it remains unclear whether it applies to specific types of memory. Here, we mapped connectivity of stimulation sites, lesions, and atrophy patterns associated with different memory types. We included 544 individuals across three datasets: transcranial magnetic stimulation (*N*=262), penetrating head trauma (*N*=169), and ischemic stroke (*N*=113). We identified a network preferentially connected to lesions and stimulation sites specifically associated with changes in visual memory. Of note, the direction of this effect was inverted depending on whether lesions or stimulation occurred at younger age or an older age, consistent with prior results. This age effect was replicated in an independent dataset of patients with preclinical Alzheimer’s disease (N=1240). To examine neuromodulation targets, we computed electrical field models for potential TMS sites that overlap with the networks derived from each stimulation or lesion dataset; the resulting targets intersected with established targets that demonstrated efficacy for treating memory impairment - precuneus, cortical-hippocampal network, and dorsolateral prefrontal cortex – with peak intersection at medial posterior parietal lobe, angular gyrus, and left anterior middle frontal gyrus, respectively. Future head-to-head clinical trials are needed to systematically compare these proposed neuromodulation targets against each other.

**One Sentence Summary:** Neuromodulation targets for visual memory diverge by age at the time of injury or stimulation.

## INTRODUCTION

Memory impairment is a symptom of a wide range of neuropsychiatric conditions. While existing treatments moderately decelerate progression of memory decline through behavioral*(1)* and pharmacological interventions*(2–4)*, brain stimulation is proposed as a complementary treatment approach. Of note, efficacy of brain stimulation depends on identifying a suitable target *(5, 6)*. For memory ability, hippocampus may seem like an ideal target as it has been classically associated with mnemonic functions*(7, 8)*. However, subcortical structures are too deep to readily access with non-invasive transcranial magnetic stimulation (TMS). While other non-invasive techniques such as focused ultrasound can potentially target hippocampus directly *(9)*, in disorders of atrophy or lesions, direct stimulation of atrophic or necrotic tissues may be less therapeutically effective. At least for these reasons, cortical targets are required to develop therapeutic protocols for non-invasive brain stimulation (NIBS).

Notably, at least three NIBS targets have shown preliminary efficacy in randomized controlled trials for treating memory decline in Alzheimer’s disease (AD) spectrum: precuneus*(10–12)*, lateral parietal lobe or the ‘cortical-hippocampal network’ *(13)*, and dorsolateral prefrontal cortex (DLPFC)*(14)*. Both the precuneus and lateral parietal targets have been selected based on their functional connectivity to hippocampus*(13)*, and other regions in the default mode network that contribute to memory phenotype. For dorsolateral prefrontal cortex, it is commonly targeted due to its involvement in executive function and symptoms of depression*(14, 15)* that may have a secondary effect on memory test scores *(14)*. While these targets are promising, it remains unclear which of the three targets should be used in large multi-site clinical trials. There is a need for an empirical approach to determine non-invasive neuromodulation targets.

Here, we draw on normative brain mapping to create a network encompassing stimulation sites that increase visual memory test scores, and lesions that decrease the same type of task performance *(16)*. This brain mapping approach has been shown to yield more effective stimulation targets across multiple neuropsychiatric disorders in retrospective studies*(17)*, and has recently been demonstrated prospectively in dystonia*(18)* and anxious depression*(19)*. We used the brain mapping technique to identify neuromodulation targets for modifying memory ability.

## RESULTS

### Normative Mapping for Stimulation Sites

We analyzed raw data from 13 transcranial magnetic stimulation (TMS) studies that used individualized cortical-hippocampal targets in healthy young adults (*n* = 262; summarized in Table 1). First, we localized the stimulation sites on a standard brain template, estimated its connectivity using a normative connectome database, and compared whole-brain stimulation site connectivity to memory scores using voxel-wise Pearson correlations, as in previous work (Fig. 1)*(16, 20)*. This analysis yielded a memory network for each study (Supplementary Fig. 1). Next, we evaluated whether the networks were similar to each other or heterogeneous between studies by computing spatial correlations that randomly permuted memory test scores between participants in each study. The memory networks were highly heterogeneous between studies (I^2^ = 87.254%, *Q* = 604.104, *mean r* = -0.05, *SD* = 0.586), characterized by a wide range of correlation values (*r* = -0.929 to 0.894), suggesting presence of possible sub-types in the TMS dataset.

**Table 1.**
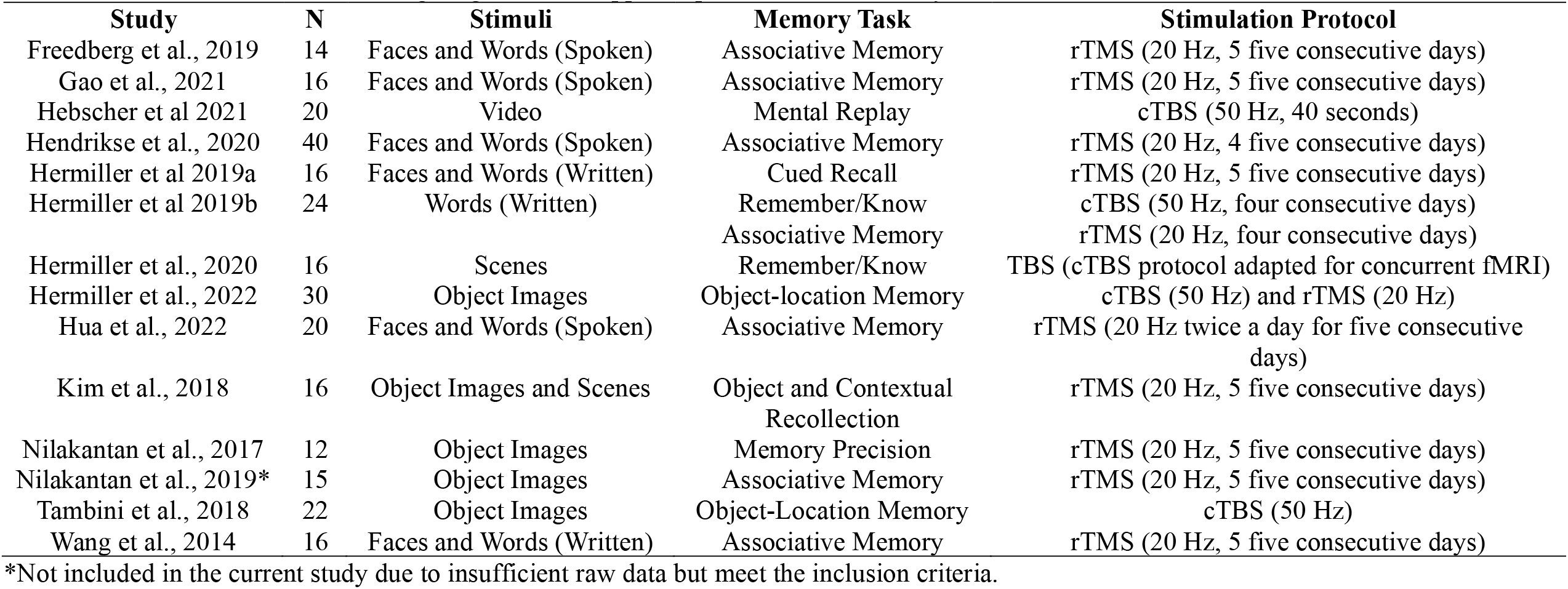
Brain Stimulation Studies Targeting Cortical-Hippocampal Network in Healthy Individuals.

| **Study** | **N** | **Stimuli** | **Memory Task** | **Stimulation Protocol** |
| --- | --- | --- | --- | --- |
| Freedberg et al., 2019 | 14 | Faces and Words (Spoken) | Associative Memory | rTMS (20 Hz, 5 five consecutive days) |
| Gao et al., 2021 | 16 | Faces and Words (Spoken) | Associative Memory | rTMS (20 Hz, 5 five consecutive days) |
| Hebscher et al 2021 | 20 | Video | Mental Replay | cTBS (50 Hz, 40 seconds) |
| Hendrikse et al., 2020 | 40 | Faces and Words (Spoken) | Associative Memory | rTMS (20 Hz, 4 five consecutive days) |
| Hermiller et al 2019a | 16 | Faces and Words (Written) | Cued Recall | rTMS (20 Hz, 5 five consecutive days) |
| Hermiller et al 2019b | 24 | Words (Written) | Remember/Know<br>Associative Memory | cTBS (50 Hz, four consecutive days)<br>rTMS (20 Hz, four consecutive days) |
| Hermiller et al., 2020 | 16 | Scenes | Remember/Know | TBS (cTBS protocol adapted for concurrent fMRI) |
| Hermiller et al., 2022 | 30 | Object Images | Object-location Memory | cTBS (50 Hz) and rTMS (20 Hz) |
| Hua et al., 2022 | 20 | Faces and Words (Spoken) | Associative Memory | rTMS (20 Hz twice a day for five consecutive days) |
| Kim et al., 2018 | 16 | Object Images and Scenes | Object and Contextual<br>Recollection | rTMS (20 Hz, 5 five consecutive days) |
| Nilakantan et al., 2017 | 12 | Object Images | Memory Precision | rTMS (20 Hz, 5 five consecutive days) |
| Nilakantan et al., 2019* | 15 | Object Images | Associative Memory | rTMS (20 Hz, 5 five consecutive days) |
| Tambini et al., 2018 | 22 | Object Images | Object-Location Memory | cTBS (50 Hz) |
| Wang et al., 2014 | 16 | Faces and Words (Written) | Associative Memory | rTMS (20 Hz, 5 five consecutive days) |

\*Not included in the current study due to insufficient raw data but meet the inclusion criteria.

**Fig. 1.**
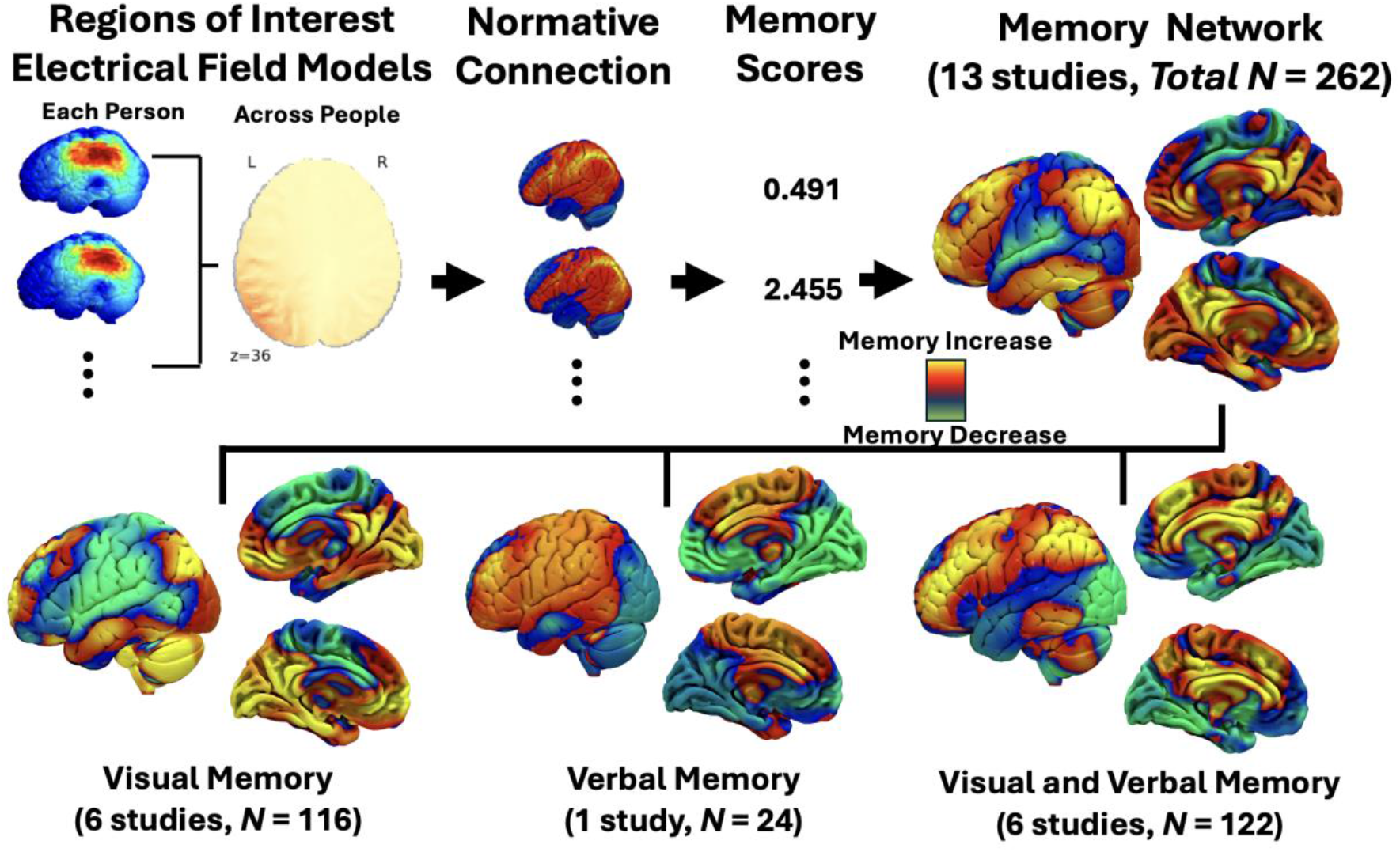
Normative Stimulation Site Mapping. Healthy young adults had received transcranial magnetic stimulation (TMS) at a lateral parietal target that was connected to hippocampus (*N* = 262). Electric field simulation models shown on the top left served as a seed to estimate normative connectivity to the rest of the brain voxels (hot colors represent higher e-field values). This procedure yielded connectivity to each stimulation site (hot colors represent higher connectivity). Correlation between the normative connectivity data and changes in memory scores before and after cortical-hippocampal target stimulation yielded a normative memory network for each study. Between memory task-specific sub-types, stimulation sites associated with improved visual verses verbal memory localized to different networks.

To probe whether this heterogeneity was driven by small sample sizes in each study, we tested whether heterogeneity is reduced when combining studies. First, we randomly split the 6 visual memory studies into 3 separate pairs across 10,000 random iterations, and recreated visual memory networks for each pair by averaging them and weighting by respective sample size. Next, we computed spatial correlations between the pairs. Increasing the sample size for each map reduced heterogeneity for studies of visual memory (I^2^ = 61.3%, *Q* = 10.56), but not for studies of combined visual/verbal outcomes (I^2^ = 85.6%, *Q* = 37.2) or when pooling all studies (I^2^ = 89.6%, *Q* = 160.8).

Thus, we generated distinct memory networks for visual memory alone versus combined visual/verbal memory and verbal memory alone. The visual memory network and the verbal memory network were inverted relative to each other (*r* = -0.778, *p* = 0.093), although this trend may be driven by small sample size in the verbal memory-only study. There was no trend towards similar networks between the visual memory and mixed visuo-verbal memory (*r* = -0.292, *p* = 0.64), and there was a weak trend towards similar networks for the verbal and visuo-verbal types (*r* = 0.569, *p* = 0.385). There was also no significant similarity or difference between circuits derived from a 20 Hz TMS protocol (*N* = 180) those derived from continuous theta burst stimulation (cTBS; *N* = 82) associated with inhibitory effects. Direct comparison between the 20 Hz and cTBS protocol-derived networks revealed no significant spatial correlation (*r* = 0.216, *p* = 0.704).

### Mapping Stimulation Sites and Penetrating Head Trauma Revealed Similar Networks

The TMS dataset alone was not sufficient to address the question of whether the network was reproducible for visual or verbal memory types. Based on previous causal brain mapping studies *(16, 21)*, we hypothesized that the TMS-derived normative networks will resemble lesion-derived networks. First, we recreated the normative network for the TMS dataset and adopted the same approach for the Traumatic Brain Injury (TBI) group with penetrating head trauma in the independent Vietnam Head Injury Study (VHIS; *N* = 169; Fig 2A)*(22)*. All Veterans were younger adults at the times of penetrating head trauma, and their cognitive tests were administered more than 15 years after the incident.

**Fig. 2.**
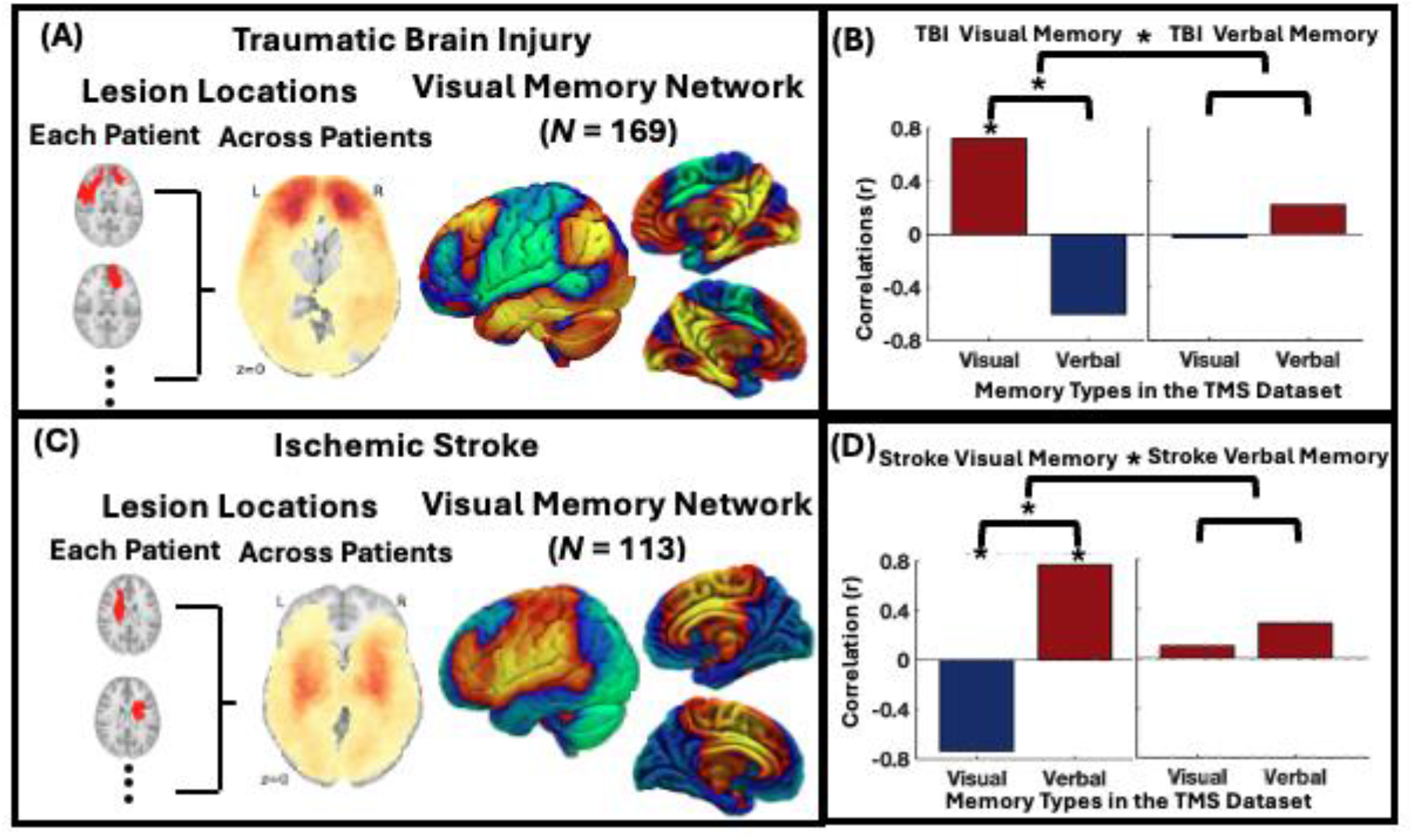
Normative Mapping Across Lesions and Stimulation Sites. Section A shows lesion locations in the traumatic brain injury (*N* = 169). Each lesion location served as a seed to estimate normative connectivity to the rest of the brain voxels. Correlation between the normative connectivity data and memory test scores yielded a visual memory network. Hot colors in the network represent regions where lesions associated with greater memory impairment. Section B shows spatial correlation between visual verses verbal memory networks from the traumatic brain injury and transcranial magnetic stimulation (TMS) datasets. Visual memory networks were specifically similar to each other, compared against verbal memory type. Section C shows lesion mapping for the stroke dataset (*N* = 113). Section D shows that visual memory networks were specifically associated with each other across the stroke and TMS datasets, but the direction of effects were inverted relative to the results for TMS and TBI datasets.

Across all TMS studies and the visual memory network derived from the TBI dataset, spatial correlation permutation analyses revealed that memory networks resembled each other (*N* = 431, *r* = 0.745, *p* = 0.002), suggesting a common network associated with improved memory after stimulation and impaired memory after a lesion. This spatial correlation remained when repeating the analysis for visual memory across both the TMS and TBI datasets (*N* = 281, *r* = 0.7403, *p* = 0.003), and after including verbal memory in the TBI dataset as a covariate (*r* = 0.718, *p* = 0.009), reaffirming that the observed similarities between the TMS and TBI datasets was specific to visual memory type. For TMS-derived verbal memory network compared against TBI-derived visual memory, spatial correlation analysis yielded a trend toward inverted networks (*N* = 193, *r* = -0.601, *p* = 0.078). For other pairs of correlations, no statistically significant results emerged (*r* < 0.225, *p* > 0.58).

Given the diverging results for visual or verbal memory types, we assessed a possible interaction between two factors: memory type (visual or verbal), and dataset (TMS or TBI). Comparing spatial correlations between the factors revealed a significant interaction (*absolute average difference r* = 0.834, *permuted p* = 0.019), suggesting that visual and verbal memory mapped to significantly different networks across TMS and TBI datasets (Fig 2B). Follow-up pair-wise analyses revealed that the interaction was primarily driven by the similarity between visual memory networks across the TMS and TBI datasets, compared against inverted TMS-derived verbal and TBI-derived visual memory networks (*absolute average difference r* = 1.342, *p* < 0.001). Other pair-wise comparisons revealed no significant difference (*absolute average difference r* < 0.858, *p* > 0.081).

### Mapping Stimulation Sites and Acute Lesions Following Ischemic Stroke Revealed Inverted Networks

We examined reproducibility of the observed similarity between visual memory networks across stimulation and lesion datasets with an independent stroke dataset from the Washington University Stroke Study (*N* = 113; Fig 2C). Most patients in the stroke dataset were older adults (*Mean Age* = 53.646, *SD* = 11.021). We recreated the TMS-derived memory network and adopted the same normative mapping approach for the stroke dataset. We expected that memory networks would resemble each other across the TMS and stroke datasets, in line with our previous spatial correlations analyses across stimulations and lesions. However, memory network across all TMS studies and the stroke-derived visual memory network, the spatial correlation analysis revealed a trend toward inverted networks (*N* = 375, spatial *r* = -0.509, *p* = 0.098), suggesting that the networks were associated with less improvement in memory task performance following stimulation and memory impairment following lesions. This result was strengthened when repeating the analysis for the visual memory type across both the TMS and stroke datasets (*r* = -0.745, *p* = 0.002), and after controlling for verbal memory scores in the stroke dataset (*r* = -0.744, *p* = 0.003), substantiating that the results were specific to visual memory type. For a TMS-derived verbal memory and Stroke-derived visual memory types, the networks were significantly similar to each other (*r* = 0.774, *p* < 0.001). For other pairs of correlations, no statistically significant results emerged (*r* < 0.283, *p* > 0.41).

To investigate a potential interaction, we conducted a 2 x 2 factorial analysis for memory type (visual or verbal), and dataset (TMS or stroke). This analysis revealed a significant interaction (*absolute average difference r* = 0.83, *permuted p* = 0.016; Fig 2D). Follow-up pair-wise analyses revealed that visual memory networks were inverted between the TMS and stroke datasets, compared against the TMS-derived verbal memory network and Stroke-derived visual memory network (*absolute average difference r* = 1.342, *p* < 0.001). Other pair-wise comparisons revealed no significant difference (*absolute average difference r* < 0.837, *p* > 0.066).

### Age Varied Between Groups

Given that the TBI and stroke lesion datasets yielded diverging results, we investigated whether effects of lesions were different for different groups. Based on previous work suggesting that the effects of stimulation on cognition may become inverted as a function of age and neurodegeneration*(23)*, we hypothesized that lesion-derived networks might be inverted between age groups: the TBI group sustained brain injury as young adults, while the stroke group was more likely to be affected later in life. Spatial correlation analysis for TBI and Stroke datasets revealed that the visual memory networks were inverted (r = -0.642, p = 0.047).

Next, we investigated how the stroke-derived networks compare against an independent group of older patients (*N* = 1240, *mean age* = 72, *SD* = 4.485; *Male:Female ratio* = 509:731) with positive amyloid beta PET scans that suggested the presence of preclinical Alzheimer’s disease (AD)*(24)*. We postulated that datasets involving older adults – preclinical AD and stroke – would yield similar visual memory networks, as opposed to the TBI dataset where patients sustained a head injury as younger adults. For the preclinical AD and Stroke datasets, visual memory networks were similar across the datasets (*n* = 1353, spatial *r* = 0.421, *p* = 0.288) only after including age as a covariate marginally strengthened the spatial correlation (spatial *r* = 0.644, *p* = 0.055, one-tailed *p* = 0.028). For the preclinical AD and TBI datasets, visual memory networks were inverted albeit not statistically significant both before and after controlling for age (*n* = 1418, spatial *r* = -0.366, *p* = 0.395; spatial *r* = -0.479, *p* = 0.455).

Next, we examined whether age was a covariate of visual memory scores for each dataset. Whereas patients with TBI who were older at the time of phenotyping trended toward having higher memory scores (*r* = 0.137, *p* = 0.075), older patients in the both the Stroke (*r* = -0.277, *p* = 0.003) and preclinical AD datasets (*r* = -0.257, *p* < 0.001) had lower memory scores. Based on these results, we directly compared the observed correlation for the Veterans in the TBI dataset against older adults in the stroke and preclinical AD datasets. Both the stroke-TBI (z = 3.431, p < 0.001) and preclinical AD-TBI (*z* = 4.835, *p* < 0.001) comparisons revealed that age was a significantly different covariate between different groups of patients.

### Non-Invasive Brain Stimulation Targeting Revealed TMS Targets That Intersect Existing Ones

To identify potential TMS targets, we developed an empirical analysis approach that estimated overlap between stimulation electrical-field models and visual memory networks. First, we created e-field simulations for candidate TMS coil locations on the scalp: 187 standard locations in the dense 10-20 electrode placements system *(25–28)*, excluding electrode positions below the nasion or ears. The e-field models were restricted to cortical surface regions by applying the top 0.01% threshold *(29)* for each of the normalized electric field models. Next, we applied the e-field model masks to the visual memory networks derived from each dataset and computed maximum spatial correlation test between datasets for all potential TMS coil locations.

Across the stimulation and lesion datasets (i.e., TMS, TBI, and Stroke), the site of greatest absolute maximum spatial correlation was at medial superior parietal part of the scalp (CPP1h; MNI: -14, -79, 83; *maximum absolute r* = 0.959, *95*^*th*^ *percentile* = 0.95, *permuted p* = 0.025; see Fig3A) overlying superior parietal lobe (MNI: -14, -67, 64). The electrical field model for this superior parietal target intersected with an e-field model of a stimulation site designed to target the precuneus (MNI: 0, -65, 45) that has previously demonstrated efficacy for decelerating progression of memory decline in Alzheimer’s Disease *(12)*. Intersection of the two targets was at centered at medial posterior parietal lobe (MNI: -9, -78, 57).

The observed posterior parietal target may have been driven by a specific pair of datasets, and different pairs may yield slightly different targets. Based on this rationale, we repeated the analysis for each pair of stimulation and lesion datasets. The TMS and Stroke datasets yielded significant sites (*maximum absolute r* = 0.99, *95*^*th*^ *percentile* = 0.979, *permuted p* = 0.01) at CP3 (MNI: -66, -57, 61), and CP4 (MNI: 69, -58, 54) overlying bilateral parietal lobes (MNI: -53, -49, 48; 54, -54, 44; see Fig 3B). Of note, the observed spatial correlation was negative, suggesting that the effect of TMS for healthy volunteeers and and acute lesions follwing stroke on visual memory task performance were opposite. The left lateral parietal target intersected with the cortical-hippocampal target (MNI: -50, -65, 33) that previously demonstrated efficacy for slowing cognitive decline in AD *(30)*. The center of intersection between the left lateral parietal and cortical-hippocampal targets was at angular gyrus (MNI: -56, -59, 41). For other pairs of stimulation or lesion datasets, spatial correlation analysis revealed no significant targets (*maximum absolute r* < 0.871, *95*^*th*^ *percentile* > 0.876, *permuted p* > 0.062).

**Fig. 3.**
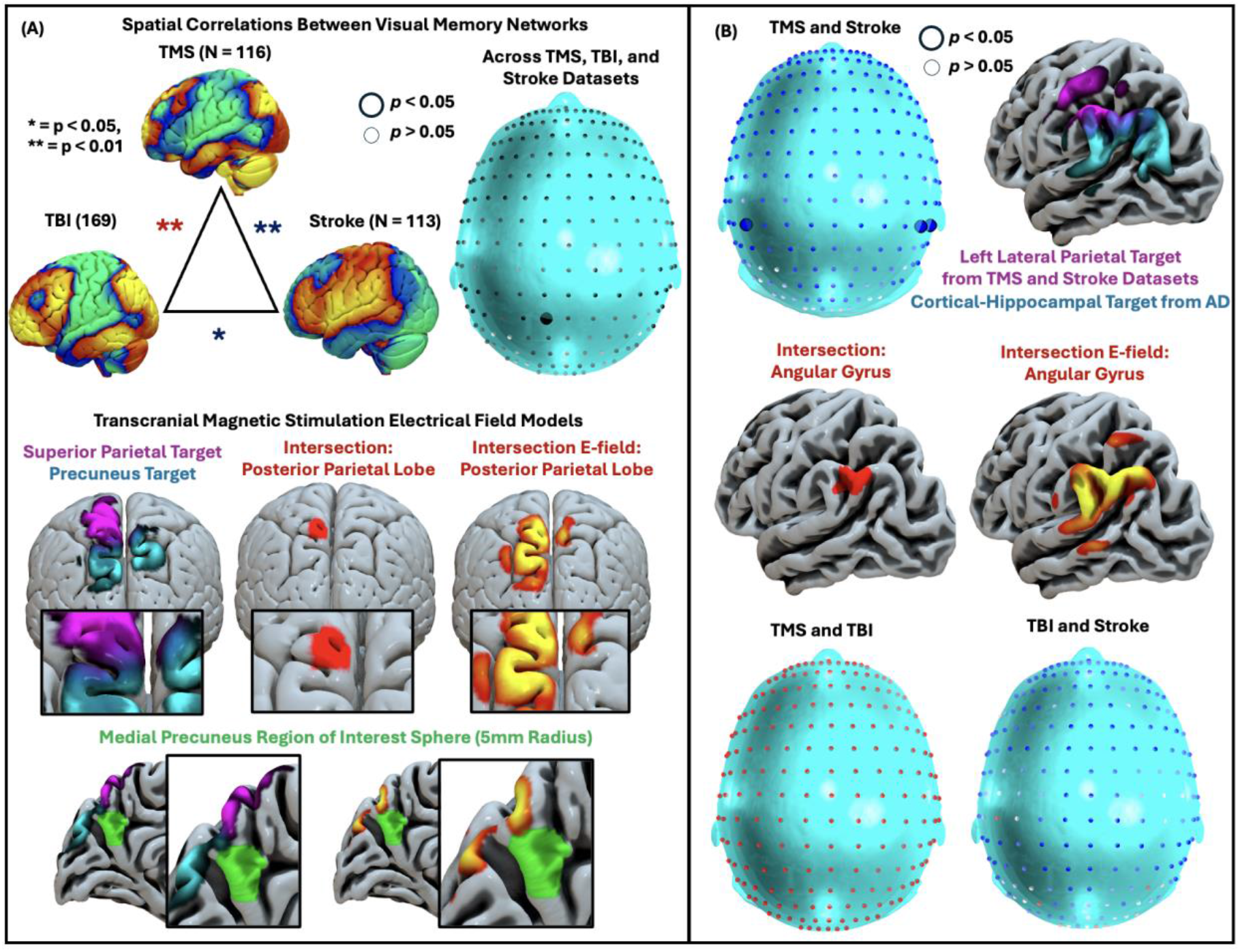
Transcranial Magnetic Stimulation Targeting Analysis Revealed Potential Targets at Medial Posterior Parietal Lobe, and Angular Gyrus. (A)Visual memory networks were spatially correlated with each other across the transcranial magnetic stimulation (TMS), traumatic brain injury (TBI), and stroke datasets. We repeated this analysis for cortical regions by creating TMS electrical field simulation models, and restricting the analysis to regions that fall within the top 0.01% of the e-field models. Potential stimulation sites on the scalp were 187 locations in the extended 10-20 placement system, excluding locations below the nasion or ears. This targeting analysis revealed that the site with peak correlation (two-tailed) was overlying medial superior parietal lobe. While the larger circle represent a statistically significant site (*p* < 0.05), smaller circles indicate no statistical significance. The medial superior parietal lobe intersected the established precuneus target that previously demonstrated efficacy for treating memory impairment*(12)*. The center of the intersection was at medial posterior parietal lobe (MNI: -9, -78, 57). (B) For each pairs of datasets, the TMS and stroke datasets yielded a lateral parietal target that intersected a cortical-hippocampal target derived from patients with Alzheimer’s disease (AD) and demonstrated efficacy for treating cognitive decline in AD*(30)*. Other pairs of datasets did not reveal statistically significant stimulation sites (indicated by small circles for all potential sites).

We also repeated the targeting analysis after adding the preclinical AD atrophy dataset. The atrophy-based mapping may yield weaker causal inference than lesion-based mapping*(21, 31, 32)*, has not been validated for TMS target localization, and was not part of our *a priori* hypothesis, so this analysis was treated as exploratory. Across all stimulation, lesion, and preclinical AD datasets, TMS targeting analysis revealed no statistically significnat site (*maximum absolute r* = 0.918, *95*^*th*^ *percentile* = 0.933, *permuted p* = 0.99; Fig 4A). Given the large sample size in the preclinical AD datset (*N* =1240), we reasoned that including the preclinical AD dataset in the analysis may haved obscured significant results from the causal datasets (i.e., TMS, TBI, and stroke). To explore whether the preclinical AD dataset poitned to a distinct target, we estimated overlap between stimulation e-field models and the respective visual memory network. This analysis revealed multiple sites across the scalp (*maximum absolute overlap score* = 18.662, *95*^*th*^ *percentile* = 11.821, *p* < 0.01; Fig 4B). To narrow-down on a relative peak, we identified sites that replicated across both e-field models with and without the top 0.01% threshold. This analysis reveaeld a relative peak in the left frontal part of the scalp (F7h; MNI: -66, 42, 6; Fig 4B), overlying inferior frontal gyrus (MNI: -53, 32, 6). The e-field model for the inferior frontal gyrus target intersected the ‘Beam F3’ dorsolateral prefrontal cortex (DLPFC) target (MNI: -38, 44, 26) previously used to increase neuropsychological test scores in AD*(33–36)*. The center of intersection between the inferior frontal gyrus and the Beam F3 target was at the anterior middle frontal gyrus part of DLPFC (MNI: -41, 47, 15). Conversely, we found a relatively negative peak at an occipital location on the scalp (POO1h MNI: 7, -118, 7) overlying occipital pole (MNI: 11, -103, 7). While this occipital target did not overlap with any of the existing targets (i.e., precuneus, lateral parietal, or DLPFC), the negative voxels extended to a layer of cortical surface regions between the medial posterior parietal target and positive voxels in the mesial precuneus region of interest.

**Fig. 4.**
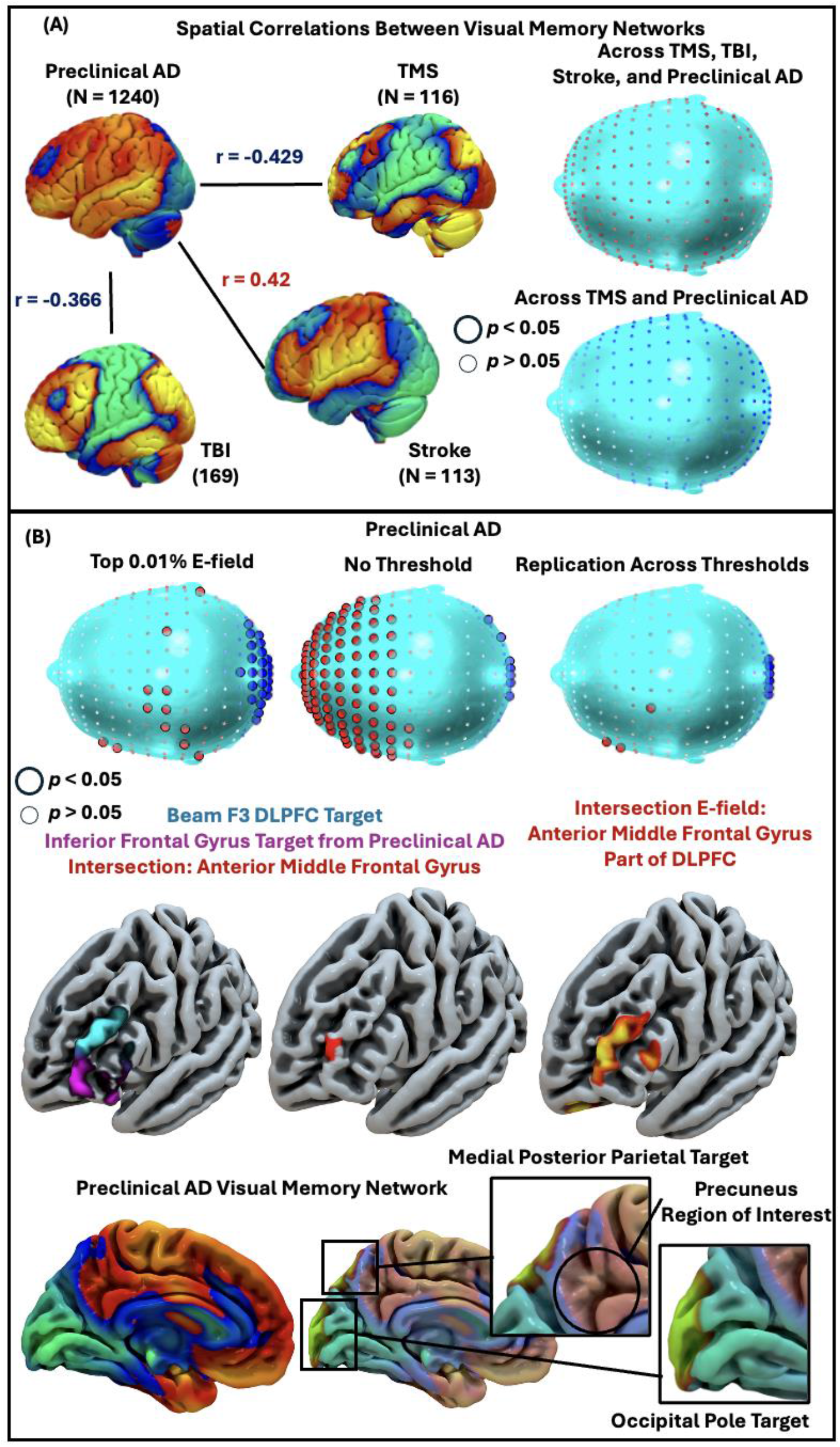
Transcranial Magnetic Stimulation Targeting Analysis for Preclinical Alzheimer’s Disease Reveals a Distinct Target in Middle Frontal Gyrus of Dorsolateral Prefrontal Cortex. Normative mapping for atrophy patterns in preclinical Alzheimer’s disease (AD) revealed a corresponding visual memory network (A). When comparing the preclinical AD-derived network to networks derived from the stimulation or lesion datasets, spatial correlation was weaker than what was observed between just the causal datasets. Given the large sample size in the preclinical AD dataset (N=1240), we reasoned that this dataset may point to a distinct network target from the causal datasets. Neuromodulation targeting analysis for preclinical AD revealed a relative peak at inferior frontal gyrus that intersected the standard Beam F3 target overlying dorsolateral prefrontal cortex (DLPFC) that previously demonstrated efficacy for treating cognitive decline in AD *(35, 36)*. The center of intersection between the two targets was at anterior middle frontal gyrus part of DLPFC (MNI: -41, 47, 15). Conversely, we found a relatively negative peak at occipital pole; these negative voxels extended to a layer of cortical surface regions between the medial posterior parietal target and positive voxels in mesial precuneus.

## DISCUSSION

We created memory-related brain maps based on normative functional connectivity of stimulation sites and lesions that modify visual memory test scores. This yielded several important implications when designing brain stimulation protocols for memory. First, visual memory mapped to a specific circuit target, suggesting that it should be measured independently of other cognitive variables such as verbal memory in brain stimulation studies. Second, effects of stimulation and lesions converged in some cases and diverged in other cases; the direction of this effect may differ depending on age of the lesion or age of the patient, potentially due to background neurodegeneration or long-term compensatory effects. Third, the proposed neuromodulation targets align with results of prior studies that have shown promising effects of TMS for memory impairment: precuneus *(10, 11)*, lateral parietal lobe*(37)*, and dorsolateral prefrontal cortex*(14)*.

### Brain Mapping

Our central hypothesis was that stimulation sites associated with greater memory improvement would map onto a common normative brain map. We focused on specificity of different memory types, as global memory have previously been localized using similar methods*(38, 39)*. Visual memory localized to different patterns of circuitry compared against verbal memory, implying that different targets may be necessary for different symptom clusters. Extending this to lesion datasets, visual memory networks were specifically associated with each other, compared against networks involving a verbal memory type. These findings are in line with previous brain mapping studies that found symptom-specific circuit targets for other disorders, and distinct stimulation targets for each symptom *(40, 41)*. In the case of depression and anxiety, prospective validation of the symptom-specific targets revealed symptom-specific remission *(19)*. These findings underscore the importance of examining specificity of neuromodulation targets for future clinical trials.

We also hypothesized that stimulation sites and lesions associated with visual memory task performance would map onto a common brain map. We found diverging effects – whereas the TMS dataset and the Vietnam Head Injury Study (VHIS) cohort yielded similar visual memory networks, their connectivity profiles were inverted compared against the network derived from the Washington University Stroke cohort. There are at least two possible explanations for this finding. One explanation is that stimulation and lesions induce opposite effects: whereas stimulation to a network can improve a symptom, lesions to the same network will induce the same symptom. In this case, TBI and TMS converged in the expected direction, while stroke lesions yielded inverted effects. This may be driven by the fact that stroke patients were older at the time of the lesion and had underlying cerebrovascular disease, consistent with prior literature suggesting that the directional effects of stimulation can become inverted by the presence of underlying neurodegeneration *(23)*. This explanation is further supported by the fact that atrophy in preclinical Alzheimer’s disease localized to the same visual memory network as stroke. The alternative explanation is that stimulation and lesions both disrupt the circuit, particularly in cases where the circuit is otherwise functioning normally. In this case, stroke and TMS converged in the expected direction, while TBI lesions yielded inverted effects. This may be driven by the long 15-year lag between injury and phenotyping, during which extensive rehabilitative and compensatory remodeling may have occurred *(42)*. Implications of these age-related differences extend to NIBS treatments for earlier and later stages of disease progression; for instance, stimulation of the lateral parietal-hippocampal network tends to reproducibly improve memory in young healthy volunteers, but yielded mixed results as age of participants varied*(13, 43–59)*; In a wider age range (i.e., 18-55 years), stimulation of lateral parietal lobe yielded no significant effect on associative memory*(46)*, direct comparison between younger verses older adults showed that the effect of stimulation was only in younger adults*(50)*, and an effect of stimulation on immediate but not short-term or long-term memory in AD*(59)*. Based on these findings, future clinical trials of any given TMS target may benefit from minimizing heterogeneity in age.

### Non-invasive Neuromodulation Targeting

We examined three existing TMS targets that demonstrated preliminary efficacy for treating memory impairment, a key symptom of Alzheimer’s Disease: precuneus*(10, 12)*, lateral parietal lobe*(13)*, and DLPFC*(14)*. Based on analyses for TMS sites, lesions, and atrophy, we identified a targeting approach that refine the targets for future clinical trials.

We identified a location in the medial posterior parietal lobe where the precuneus target intersected the superior parietal location with the strongest relative spatial correlation across the stimulation and lesion datasets. It is plausible that the documented efficacy of the precuneus target was partly driven by incidental intersection with this posterior parietal lobe target. Precuneus has conventionally been thought to be part of the default mode network (DMN)*(10, 11)*, and this DMN-like precuneus region of interest was part of the visual memory network derived from the independent preclinical AD dataset. Of note, we found a layer of negative cortical surface voxels between the precuneus region of interest and the posterior parietal target TMS coil location on the scalp. Since TMS cannot directly reach deep regions without first penetrating superficial regions, the reported clinical effects of precuneus stimulation may be driven by the overlying negative cluster. This characteristic for the precuneus target is distinguishable from both the lateral parietal and DLPFC targets, as these two targets directly map onto cortical surface regions with positive voxels derived from patients with preclinical AD. As it remains unclear whether positive or negative voxels in this type of map constitute better targets, a prospective trial that directly compares the NIBS targets is warranted.

We also found a separate target at angular gyrus that intersected the cortical-hippocampal target from a seminal previous trial for AD spectrum*(30)*, and the posterior supramarginal gyrus target derived from the TMS and Stroke datasets. These lateral parietal target space demonstrated preliminary efficacy in previous TMS studies in health and AD *(30, 57–59)*. Despite its potential, lateral parietal targets have been underused in randomized controlled trials for AD – previous trials have often adopted a DLPFC or a precuneus target *(14)*. There may be benefit to additional trials including the angular gyrus target in a systematic trial.

Our proposed left DLPFC target space is broadly consistent with previous randomized controlled trials for AD*(35, 60–75)*, although our precise middle frontal gyrus target MNI coordinate (−41, 47, 15) is relatively lateral to the standard Beam F3 target *(33–35)*. These RCTs demonstrated efficacy of the DLPFC target space for decelerating progression of cognitive/memory decline in AD*(14)*, although these results can be challenging to interpret given that the DLPFC target has been classically used to treat other neuropsychiatric conditions such as depression*(15)*. It is plausible that stimulation of DLPFC affected comorbid depression, compensation strategies involving executive functions that draw on DLPFC, and other cognitive abilities that contribute to overall memory phenotype*(14)*. For example, DLPFC is associated with a range of cognitive abilities including selective retrieval that contribute to memory task performance*(76, 77)*. Therefore, effects of DLPFC stimulation on memory phenotype should be tested controlling for symptoms of depression and executive function.

### Limitations

The current retrospective data analysis has limitations. There is no prospective validation of our proposed neuromodulation targets, thus providing only indirect evidence for efficacy based on their overlap with previous targets. It also limits inference about collective and relative efficacy of these targets compared against each other. For instance, it remains unclear whether all three or one of these target stimulations should be administered to patients with memory impairment. Future studies may benefit from systematically comparing individualized TMS site connectivity to our three targets (i.e., medial posterior parietal lobe, left angular gyrus, and anterior middle frontal gyrus), which we were unable to do because most randomized controlled trials for memory disorders did not measure variability in individual TMS sites. We also had no empirical control over participants’ demographic and neuropsychiatric profiles, which is particularly notable because age and depression/anxiety can both covary with memory test scores *(78)* and can both mediate the effects of stimulation*(79, 80)*. Clinical trials of TMS for mnemonic function may benefit from tighter control of age, neurodegeneration, and psychiatric comorbidities as an inclusion criterion or a covariate. We also could not conclusively characterize the nature of the relative inversion between memory localization in TBI versus stroke lesions. This raises questions about whether positive or negative parts of the circuit should be treated as potential stimulation targets, so target-specific clinical trials are necessary before translating these findings into clinical practice.

## Conclusions

Our targeting approach may help refine existing correlational or scalp-based methods. Causal methods with precise localization have previously been shown to improve brain stimulation targeting across multiple disorders. However, future studies are still needed to validate these targets prospectively. To facilitate this process, we provide MNI coordinates and scalp approximations for the three key targets: medial posterior parietal lobe, left angular gyrus, and anterior middle frontal gyrus part of the DLPFC. While stimulation of all three targets might prove to be most effective, it is possible that different patients respond to different targets, one of the targets systematically outperform others, or none of them show efficacy controlling for age, symptoms of depression, or executive function. Such a prospective trial will provide insight about which targets to use in larger multi-site clinical trials, a crucial step for the emerging field of NIBS for treating memory impairment.

## MATERIALS AND METHODS

### Data

We included a total of 1784 individuals across four datasets. Dataset 1 consisted of 13 prior studies involving healthy volunteers who received transcranial magnetic stimulation (TMS) at the cortical-hippocampal network target based on its involvement in memory task performance (N = 262; Table 1). Dataset 2 consisted of patients who sustained penetrating traumatic brain injury (TBI), primarily in prefrontal cortex with some in temporal cortex, during military service in the Vietnam War (N = 169)*(22)*. The Veterans’ cognitive test scores were taken from Phase 2 of the Vietnam Head Injury Study, at least 15 years after the injury. Dataset 3 consisted of patients with acute stroke-induced lesions, primarily in subcortical regions, recruited in an observational study at Washington University in St. Louis (N = 113)*(81)*. Dataset 4 consisted of individuals who tested positive for beta-amyloid plaques and are at risk of Alzheimer’s disease, but they were asymptomatic at the time of testing (N = 1240) *(24)*. The four datasets collectively provide a wide spread of brain lesion and stimulation target locations, sufficient for comprehensive brain mapping. Although dataset 4 is larger in sample size, it was expected to yield a smaller effect size because all patients were asymptomatic and because atrophy and amyloid deposition generally show weaker causal brain mapping results than lesions or stimulation sites.

### Normative Brain Network Mapping

We adopted lesion network mapping*(20)* and more broadly, convergent causal brain mapping*(16, 21)*. Here, the brain lesion, stimulation and atrophy locations serve as seed regions that map onto a normative resting-state brain network from a large sample of healthy volunteers (*N* = 1000)*(82, 83)*. This procedure provided normative connectivity to each seed region, where positive values reflect higher functional connectivity to the seed region and negative values reflect stronger negative connectivity. Correlation between the normative connectivity data and memory scores yielded normative memory network.

### Memory Scores

For the TMS dataset, there were multiple independent studies, each with different episodic memory tasks. Across all the TMS studies, the key measure of task performance is change in memory score before and after target stimulation or, alternatively, change in memory scores following target and control stimulation. For the TBI dataset, the memory score of interest was a combined score across visual memory tasks (i.e., the Face Acquisition and Recognition task *(84)*, and the Recurring Figure test *(85)*). Respective memory test scores in the TBI dataset were normalized and then averaged for each patient. In the stroke dataset, the memory score used is the delayed recognition memory discrimination index (i.e., d-prime) in the Brief Visuospatial Memory Test*(86)*. For the atrophy dataset, we used the Free and Cued Selective Reminding Test *(87)*. To ensure that the direction of memory scores was aligned across datasets, we inversed the visuospatial memory test scores such that higher scores in the brain lesion and preclinical AD datasets reflect worse memory ability.

### Analysis Approach: Non-invasive Neuromodulation Targeting

We developed an empirical approach to identify potential non-invasive neuromodulation coil positions across the scalp. Potential sites were 187 locations in the dense 10-20 electrode placement system*(25–28)*, excluding positions below the nasion or ears. For each location, we created electric field simulation models using a simulation software (SimNIBS)*(88)*. We normalized the e-field models and applied top 0.01% threshold *(29)* to (a) restrict analyses to cortical surface regions amenable for non-invasive stimulation, as opposed to subcortical regions, and (b) minimize possible overlap between e-field models with each other. Next, we recreated normative visual memory networks and multiplied the voxel values with e-field models for each potential site. Between datasets, we conducted absolute maximum spatial correlation tests that permuted visual memory scores between participants *(89)* for each potential site. Across all potential sites, we estimated the absolute maximum spatial correlation and its permuted p-value.

---

**Supplementary Fig. 1:**
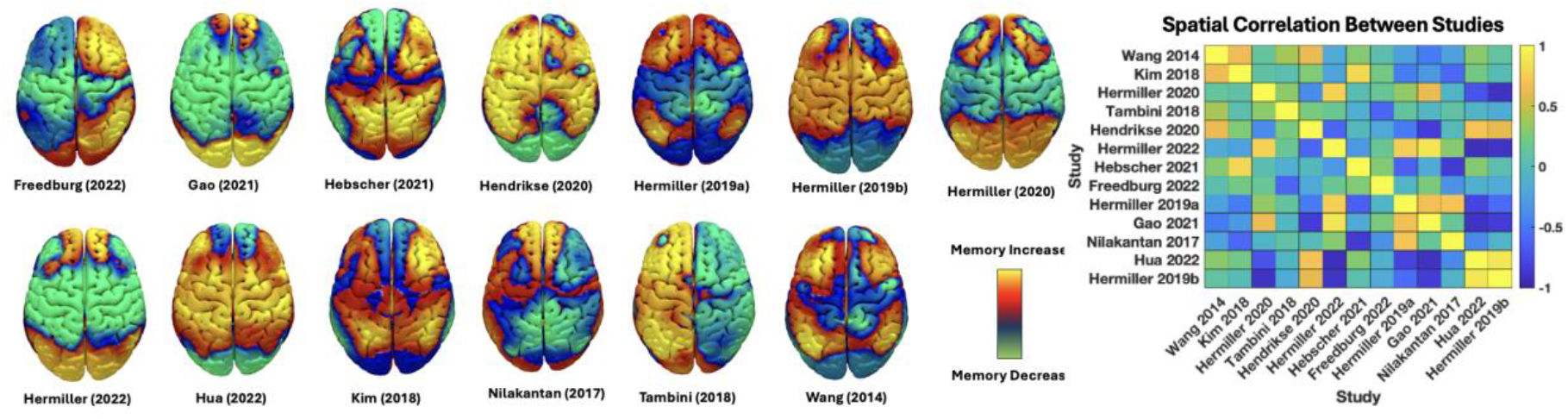
Memory Network for Each Transcranial Magnetic Stimulation Study. For each study, mapping normative connectivity to stimulation sites associated with changes in memory test scores yielded a memory network (13 studies, *Total N* = 262). Between studies, spatial correlation analyses revealed that the memory networks were heterogeneous. This variance was due to small sample sizes in each study, relative to a combination of multiple studies that provide more statistical power.

## Acknowledgments

Acknowledgments follow the references and notes but are not numbered. Start with text that acknowledges non-author contributions and then complete each of the sections below as separate paragraphs.

## Author contributions

Conceptualization: SMK, SL, SHS

Data Contribution: JSS, MF, MH, JH, MSH, GJ, AT, SC, MC, JG, JLV

Methodology: SMK, SL, NC, EY, SHS

Investigation: SMK, SL, SHS

Visualization: SMK, SHS

Funding acquisition: SHS

Project administration: SMK, SHS

Supervision: SHS

Writing – original draft: SMK, SHS

Writing – review & editing: SMK, SL, JH, GJ, AT, JG, JLV, SHS

## Data and materials availability

All data, code, and materials used in the analysis will be available upon reasonable request to any researcher for purposes of reproducing or extending the analysis.

## References and Notes

1. E. J. Lenze, M. Voegtle, J. P. Miller, B. M. Ances, D. A. Balota, D. Barch, C. A. Depp, B. S. Diniz, L. T. Eyler, E. R. Foster, T. R. Gettinger, D. Head, T. Hershey, S. Klein, J. F. Nichols, G.E. Nicol, T. Nishino, B. W. Patterson, T. L. Rodebaugh, J. Schweiger, J. S. Shimony, D. R. Sinacore, A. Z. Snyder, S. Tate, E. W. Twamley, D. Wing, G. F. Wu, L. Yang, M. D. Yingling, J. L. Wetherell, Effects of Mindfulness Training and Exercise on Cognitive Function in Older Adults: A Randomized Clinical Trial. JAMA 328, 2218 (2022).

2. C. H. van Dyck, C. J. Swanson, P. Aisen, R. J. Bateman, C. Chen, M. Gee, M. Kanekiyo, D. Li, L. Reyderman, S. Cohen, L. Froelich, S. Katayama, M. Sabbagh, B. Vellas, D. Watson, S. Dhadda, M. Irizarry, L. D. Kramer, T. Iwatsubo, Lecanemab in Early Alzheimer’s Disease. New England Journal of Medicine 388, 9–21 (2023).

3. J. R. Sims, J. A. Zimmer, C. D. Evans, M. Lu, P. Ardayfio, J. Sparks, A. M. Wessels, S. Shcherbinin, H. Wang, E. S. Monkul Nery, E. C. Collins, P. Solomon, S. Salloway, L. G. Apostolova, O. Hansson, C. Ritchie, D. A. Brooks, M. Mintun, D. M. Skovronsky, TRAILBLAZER-ALZ 2 Investigators, Donanemab in Early Symptomatic Alzheimer Disease: The TRAILBLAZER-ALZ 2 Randomized Clinical Trial. JAMA 330, 512–527 (2023).

4. S. Budd Haeberlein, P. S. Aisen, F. Barkhof, S. Chalkias, T. Chen, S. Cohen, G. Dent, O. Hansson, K. Harrison, C. von Hehn, T. Iwatsubo, C. Mallinckrodt, C. J. Mummery, K. K. Muralidharan, I. Nestorov, L. Nisenbaum, R. Rajagovindan, L. Skordos, Y. Tian, C. H. van Dyck, B. Vellas, S. Wu, Y. Zhu, A. Sandrock, Two Randomized Phase 3 Studies of Aducanumab in Early Alzheimer’s Disease. J Prev Alzheimers Dis 9, 197–210 (2022).

5. P. M. Rossini, D. Burke, R. Chen, L. G. Cohen, Z. Daskalakis, R. Di Iorio, V. Di Lazzaro, F. Ferreri, P. B. Fitzgerald, M. S. George, M. Hallett, J. P. Lefaucheur, B. Langguth, H. Matsumoto, C. Miniussi, M. A. Nitsche, A. Pascual-Leone, W. Paulus, S. Rossi, J. C. Rothwell, H. R. Siebner, Y. Ugawa, V. Walsh, U. Ziemann, Non-invasive electrical and magnetic stimulation of the brain, spinal cord, roots and peripheral nerves: Basic principles and procedures for routine clinical and research application. An updated report from an I.F.C.N. Committee. Clin Neurophysiol 126, 1071–1107 (2015).

6. A. Opitz, W. Legon, A. Rowlands, W. K. Bickel, W. Paulus, W. J. Tyler, Physiological observations validate finite element models for estimating subject-specific electric field distributions induced by transcranial magnetic stimulation of the human motor cortex. NeuroImage 81, 253–264 (2013).

7. W. B. Scoville, B. Milner, LOSS OF RECENT MEMORY AFTER BILATERAL HIPPOCAMPAL LESIONS. Journal of Neurology, Neurosurgery, and Psychiatry 20, 11 (1957).

8. E. Tulving, Episodic Memory: From Mind to Brain. Annu. Rev. Psychol. 53, 1–25 (2002).

9. A. R. Rezai, M. Ranjan, P.-F. D’Haese, M. W. Haut, J. Carpenter, U. Najib, R. I. Mehta, J. L. Chazen, Z. Zibly, J. R. Yates, S. L. Hodder, M. Kaplitt, Noninvasive hippocampal blood−brain barrier opening in Alzheimer’s disease with focused ultrasound. Proceedings of the National Academy of Sciences 117, 9180–9182 (2020).

10. T. Yokoi, H. Watanabe, H. Yamaguchi, E. Bagarinao, M. Masuda, K. Imai, A. Ogura, R. Ohdake, K. Kawabata, K. Hara, Y. Riku, S. Ishigaki, M. Katsuno, S. Miyao, K. Kato, S. Naganawa, R. Harada, N. Okamura, K. Yanai, M. Yoshida, G. Sobue, Involvement of the Precuneus/Posterior Cingulate Cortex Is Significant for the Development of Alzheimer’s Disease: A PET (THK5351, PiB) and Resting fMRI Study. Front. Aging Neurosci. 10, 304 (2018).

11. G. Koch, S. Bonnì, M. C. Pellicciari, E. P. Casula, M. Mancini, R. Esposito, V. Ponzo, S. Picazio, F. D. Lorenzo, L. Serra, C. Motta, M. Maiella, C. Marra, M. Cercignani, A. Martorana, C. Caltagirone, M. Bozzali, Transcranial magnetic stimulation of the precuneus enhances memory and neural activity in prodromal Alzheimer’s disease. NeuroImage 169, 302–311 (2018).

12. G. Koch, E. P. Casula, S. Bonnì, I. Borghi, M. Assogna, F. Di Lorenzo, R. Esposito, M. Maiella, A. D’Acunto, M. Ferraresi, L. Mencarelli, V. Pezzopane, C. Motta, E. Santarnecchi, M. Bozzali, A. Martorana, Effects of 52 weeks of precuneus rTMS in Alzheimer’s disease patients: a randomized trial. Alz Res Therapy 17, 69 (2025).

13. J. X. Wang, L. M. Rogers, E. Z. Gross, A. J. Ryals, M. E. Dokucu, K. L. Brandstatt, M. S. Hermiller, J. L. Voss, Targeted enhancement of cortical-hippocampal brain networks and associative memory. Science 345, 1054–1057 (2014).

14. S. R. Pagali, R. Kumar, A. M. LeMahieu, M. R. Basso, B. F. Boeve, P. E. Croarkin, J. R. Geske, L. C. Hassett, J. Huston, S. Kung, B. N. Lundstrom, R. C. Petersen, E. K. St. Louis, K. M. Welker, G. A. Worrell, A. Pascual-Leone, M. I. Lapid, Efficacy and safety of transcranial magnetic stimulation on cognition in mild cognitive impairment, Alzheimer’s disease, Alzheimer’s disease-related dementias, and other cognitive disorders: a systematic review and meta-analysis. International Psychogeriatrics 36, 880–928 (2024).

15. A. Pascual-Leone, B. Rubio, F. Pallardó, M.D. Catalá, Rapid-rate transcranial magnetic stimulation of left dorsolateral prefrontal cortex in drug-resistant depression. Lancet 348, 233–237 (1996).

16. S. H. Siddiqi, F. L. W. V. J. Schaper, A. Horn, J. Hsu, J. L. Padmanabhan, A. Brodtmann, R. F. H. Cash, M. Corbetta, K. S. Choi, D. D. Dougherty, N. Egorova, P. B. Fitzgerald, M. S. George, S. A. Gozzi, F. Irmen, A. A. Kuhn, K. A. Johnson, A. M. Naidech, A. Pascual-Leone, T. G. Phan, R. P. W. Rouhl, S. F. Taylor, J. L. Voss, A. Zalesky, J. H. Grafman, H. S. Mayberg, M. D. Fox, Brain stimulation and brain lesions converge on common causal circuits in neuropsychiatric disease. Nat Hum Behav 5, 1707–1716 (2021).

17. S. H. Siddiqi, M. D. Fox, Targeting Symptom-Specific Networks With Transcranial Magnetic Stimulation. Biological Psychiatry 95, 502–509 (2024).

18. Kokkonen, D. T. Corp, J. Aaltonen, J. Hirvonen, A. K. Kirjavainen, J. Rajander, J. Joutsa, Brain metabolic response to repetitive transcranial magnetic stimulation to lesion network in cervical dystonia. Brain Stimulation 17, 1171–1177 (2024).

19. J. Taylor, J. Li, C. Lin, E. Jones, S. Frandsen, C. Becker, W. Drew, D. Haj-Darwish, S. Jabbour, J. Leach, S. Palm, L. Sanderson, E. Santos, L. Sterina, S. Baratono, I. Gonsalvez, S. Lyndon, S. Snider, M. Fox, S. Siddiqi, Symptom-specific brain circuit stimulation: A head-to-head randomized trial (2024), doi:10.21203/rs.3.rs-4791291/v1.

20. M. D. Fox, Mapping Symptoms to Brain Networks with the Human Connectome. N Engl J Med 379, 2237–2245 (2018).

21. S. H. Siddiqi, K. P. Kording, J. Parvizi, M. D. Fox, Causal mapping of human brain function. Nat Rev Neurosci 23, 361–375 (2022).

22. V. Raymont, A. M. Salazar, F. Krueger, J. Grafman, “Studying Injured Minds” – The Vietnam Head Injury Study and 40 Years of Brain Injury Research. Front. Neur. 2 (2011), doi:10.3389/fneur.2011.00015.

23. C. W. Howard, M. Reich, L. Luo, N. Pacheco-Barrios, R. Alterman, A. S. Rios, M. Guo, Z. Luo, H. Friedrich, A. Pines, L. Montaser-Kouhsari, W. Drew, L. Hart, G. Meyer, N. Rajamani, M. U. Friedrich, V. Milanese, A. Lozano, for the Ad. S. R. Group, T. Picht, K. Faust, A. Horn, M. D. Fox, Cognitive outcomes of deep brain stimulation depend on age and hippocampal connectivity in Parkinson’s and Alzheimer’s disease. Alzheimer’s & Dementia 21, e70498 (2025).

24. R. A. Sperling, M. C. Donohue, R. Raman, M. S. Rafii, K. Johnson, C. L. Masters, C. H. van Dyck, T. Iwatsubo, G. A. Marshall, R. Yaari, M. Mancini, K. C. Holdridge, M. Case, J. R. Sims, P. S. Aisen, Trial of Solanezumab in Preclinical Alzheimer’s Disease. N Engl J Med 389, 1096–1107 (2023).

25. G. Deuschl, A. Eisen, Recommendations for the practice of clinical neurophysiology: guidelines of the International Federation of Clinical Neurophysiology. Electroencephalography and clinical neurophysiology. Supplement (1999) (available at https://www.semanticscholar.org/paper/Recommendations-for-the-practice-of-clinical-of-the-Deuschl-Eisen/e534ee50f10bb49a60c876050ef1d762e883da2b).

26. G. H. Klem, H. O. LuÈders, H. H. Jasper, C. Elger, The ten-twenty electrode system of the International Federation. .

27. A. Delorme, S. Makeig, EEGLAB: an open source toolbox for analysis of single-trial EEG dynamics including independent component analysis. Journal of Neuroscience Methods 134, 9–21 (2004).

28. V. Litvak, J. Mattout, S. Kiebel, C. Phillips, R. Henson, J. Kilner, G. Barnes, R. Oostenveld, J. Daunizeau, G. Flandin, W. Penny, K. Friston, EEG and MEG Data Analysis in SPM8. Computational Intelligence and Neuroscience 2011, 852961 (2011).

29. I. G. Elbau, C. J. Lynch, J. Downar, F. Vila-Rodriguez, J. D. Power, N. Solomonov, Z. J. Daskalakis, D. M. Blumberger, C. Liston, Functional Connectivity Mapping for rTMS Target Selection in Depression. AJP 180, 230–240 (2023).

30. Y. H. Jung, H. Jang, S. Park, H. J. Kim, S. W. Seo, G. B. Kim, Y.-M. Shon, S. Kim, D. L. Na, Effectiveness of Personalized Hippocampal Network–Targeted Stimulation in Alzheimer Disease: A Randomized Clinical Trial. JAMA Netw Open 7, e249220–e249220 (2024).

31. J. Taylor, C. Lin, D. Talmasov, M. A. Ferguson, F. L. Schaper, J. Jiang, M. Goodkind, J. Grafman, A. Etkin, S. H. Siddiqi, M. D. Fox, A Transdiagnostic Network for Psychiatric Illness Derived from Atrophy and Lesions. Nature human behaviour 7, 420 (2023).

32. A. T. Makhlouf, W. Drew, J. L. Stubbs, J. J. Taylor, D. Liloia, J. Grafman, D. Silbersweig, M. D. Fox, S. H. Siddiqi, Heterogeneous patterns of brain atrophy in schizophrenia localize to a common brain network. Nat. Mental Health 3, 19–30 (2025).

33. W. Beam, J. J. Borckardt, S. T. Reeves, M. S. George, An efficient and accurate new method for locating the F3 position for prefrontal TMS applications. Brain Stimul 2, 50–54 (2009).

34. A. Mir-Moghtadaei, R. Caballero, P. Fried, M. D. Fox, K. Lee, P. Giacobbe, Z. J. Daskalakis, D. M. Blumberger, J. Downar, Concordance Between BeamF3 and MRI-neuronavigated Target Sites for Repetitive Transcranial Magnetic Stimulation of the Left Dorsolateral Prefrontal Cortex. Brain Stimul 8, 965–973 (2015).

35. X. Wu, G.-J. Ji, Z. Geng, L. Wang, Y. Yan, Y. Wu, G. Xiao, L. Gao, Q. Wei, S. Zhou, L. Wei, Y. Tian, K. Wang, Accelerated intermittent theta-burst stimulation broadly ameliorates symptoms and cognition in Alzheimer’s disease: A randomized controlled trial. Brain Stimulation 15, 35–45 (2022).

36. X. Wu, Y. Yan, P. Hu, L. Wang, Y. Wu, P. Wu, Z. Geng, G. Xiao, S. Zhou, G. Ji, B. Qiu, L. Wei, Y. Tian, H. Liu, K. Wang, Effects of a periodic intermittent theta burst stimulation in Alzheimer’s disease. Gen Psychiatr 37, e101106 (2024).

37. T. H.-Y. Wang, Investigations of age-related effects on the neural correlates of recollection and familiarity. Dissertation Abstracts International: Section B: The Sciences and Engineering 74, No Pagination Specified (2014).

38. M. A. Ferguson, C. Lim, D. Cooke, R. R. Darby, O. Wu, N. S. Rost, M. Corbetta, J. Grafman, M. D. Fox, A human memory circuit derived from brain lesions causing amnesia. Nat Commun 10, 3497 (2019).

39. C. W. Howard, S. Madan, A. Garimella, F. Schaper, I. Kletenik, M. C. Ng, P. Mosley, J. Grafman, R. Bakshi, B. Glanz, L. Fosdick, A. Johnson, R. Colyer, C. G. Lyketsos, M. Morton-Dutton, J. Giftakis, Y. Temel, R. P. W. Rouhl, J. H. Ko, R. Schmahl, J. C. Baldermann, Ö. Onur, P. A.-Montemayor, V. Visser-Vandewalle, J. Kuhn, M. Corbetta, R. S. Fisher, T. Picht, K. Faust, M. Hermiller, J. Voss, T. Chitnis, M. K. Kahana, G. S. Smith, A. Lozano, S. H. Siddiqi, A. Horn, M. D. Fox, A Network Target for Memory Dysfunction Derived from Brain Lesions and Stimulations, 2026.03.10.26348082 (2026).

40. S. H. Siddiqi, S. F. Taylor, D. Cooke, A. Pascual-Leone, M. S. George, M. D. Fox, Distinct Symptom-Specific Treatment Targets for Circuit-Based Neuromodulation. AJP 177, 435–446 (2020).

41. N. Rajamani, H. Friedrich, K. Butenko, T. Dembek, F. Lange, P. Navrátil, P. Zvarova, B. Hollunder, R. M. A. de Bie, V. J. J. Odekerken, J. Volkmann, X. Xu, Z. Ling, C. Yao, P. Ritter, W.-J. Neumann, G. P. Skandalakis, S. Komaitis, A. Kalyvas, C. Koutsarnakis, G. Stranjalis, M. Barbe, V. Milanese, M. D. Fox, A. A. Kühn, E. Middlebrooks, N. Li, M. Reich, C. Neudorfer, A. Horn, Deep brain stimulation of symptom-specific networks in Parkinson’s disease. Nat Commun 15, 4662 (2024).

42. V. Raymont, J. Grafman, in Progress in Brain Research, (Elsevier, 2006), vol. 157, pp. 199–206.

43. M. Freedberg, J. A. Reeves, A. C. Toader, M. S. Hermiller, J. L. Voss, E. M. Wassermann, Persistent Enhancement of Hippocampal Network Connectivity by Parietal rTMS Is Reproducible. eNeuro 6, ENEURO.0129-19.2019 (2019).

44. X. Gao, Q. Hua, R. Du, J. Sun, T. Hu, J. Yang, B. Qiu, G.-J. Ji, K. Wang, Associative memory improvement after 5 days of magnetic stimulation: A replication experiment with active controls. Brain Research 1765, 147510 (2021).

45. M. Hebscher, J. E. Kragel, T. Kahnt, J. L. Voss, Enhanced reinstatement of naturalistic event memories due to hippocampal-network-targeted stimulation. Current Biology 31, 1428-1437.e5 (2021).

46. J. Hendrikse, J. P. Coxon, S. Thompson, C. Suo, A. Fornito, M. Yücel, N. C. Rogasch, Multi-day rTMS exerts site-specific effects on functional connectivity but does not influence associative memory performance. Cortex 132, 423–440 (2020).

47. M. S. Hermiller, E. Karp, A. S. Nilakantan, J. L. Voss, Episodic memory improvements due to noninvasive stimulation targeting the cortical–hippocampal network: A replication and extension experiment. Brain and Behavior 9, e01393 (2019).

48. M. S. Hermiller, S. VanHaerents, T. Raij, J. L. Voss, Frequency-specific noninvasive modulation of memory retrieval and its relationship with hippocampal network connectivity. Hippocampus 29, 595–609 (2019).

49. M. S. Hermiller, Y. F. Chen, T. B. Parrish, J. L. Voss, Evidence for Immediate Enhancement of Hippocampal Memory Encoding by Network-Targeted Theta-Burst Stimulation during Concurrent fMRI. J. Neurosci. 40, 7155–7168 (2020).

50. M. S. Hermiller, S. Dave, S. L. Wert, S. VanHaerents, M. Riley, S. Weintraub, M. M. Mesulam, J. L. Voss, Evidence from theta-burst stimulation that age-related de-differentiation of the hippocampal network is functional for episodic memory. Neurobiology of Aging 109, 145–157 (2022).

51. Q. Hua, Y. Zhang, Q. Li, X. Gao, R. Du, Y. Wang, Q. Zhou, T. Zhang, J. Sun, L. Zhang, G. Ji, K. Wang, Efficacy of twice-daily high-frequency repetitive transcranial magnetic stimulation on associative memory. Front. Hum. Neurosci. 16, 973298 (2022).

52. S. Kim, A. S. Nilakantan, M. S. Hermiller, R. T. Palumbo, S. VanHaerents, J. L. Voss, Selective and coherent activity increases due to stimulation indicate functional distinctions between episodic memory networks. Sci. Adv. 4, eaar2768 (2018).

53. A. S. Nilakantan, D. J. Bridge, E. P. Gagnon, S. A. VanHaerents, J. L. Voss, Stimulation of the Posterior Cortical-Hippocampal Network Enhances Precision of Memory Recollection. Current biology : CB 27, 465–470 (2017).

54. A. Tambini, D. E. Nee, M. D’Esposito, Hippocampal-targeted theta-burst stimulation enhances associative memory formation. Journal of cognitive neuroscience 30, 1452–1472 (2018).

55. A. S. Nilakantan, M.-M. Mesulam, S. Weintraub, E. L. Karp, S. VanHaerents, J. L. Voss, Network-targeted stimulation engages neurobehavioral hallmarks of age-related memory decline. Neurology 92 (2019), doi:10.1212/WNL.0000000000007502.

56. H. A. Velioglu, L. Hanoglu, Z. Bayraktaroglu, G. Toprak, E. M. Guler, M. Y. Bektay, O. Mutlu-Burnaz, B. Yulug, Left lateral parietal rTMS improves cognition and modulates resting brain connectivity in patients with Alzheimer’s disease: Possible role of BDNF and oxidative stress. Neurobiology of Learning and Memory 180, 107410 (2021).

57. Y. Hu, Y. Jia, Y. Sun, Y. Ding, Z. Huang, C. Liu, Y. Wang, Efficacy and safety of simultaneous rTMS–tDCS over bilateral angular gyrus on neuropsychiatric symptoms in patients with moderate Alzheimer’s disease: A prospective, randomized, sham-controlled pilot study. Brain Stimulation 15, 1530–1537 (2022).

58. Y. Jia, L. Xu, K. Yang, Y. Zhang, X. Lv, Z. Zhu, Z. Chen, Y. Zhu, L. Wei, X. Li, M. Qian, Y. Shen, W. Hu, W. Chen, Precision Repetitive Transcranial Magnetic Stimulation Over the Left Parietal Cortex Improves Memory in Alzheimer’s Disease: A Randomized, Double-Blind, Sham-Controlled Study. Front. Aging Neurosci. 13 (2021), doi:10.3389/fnagi.2021.693611.

59. L. Wei, Y. Zhang, J. Wang, L. Xu, K. Yang, X. Lv, Z. Zhu, Q. Gong, W. Hu, X. Li, M. Qian, Y. Shen, W. Chen, Parietal-hippocampal rTMS improves cognitive function in Alzheimer’s disease and increases dynamic functional connectivity of default mode network. Psychiatry Research 315, 114721 (2022).

60. J. Cheng, J. K. Fairchild, M. W. McNerney, A. Noda, J. W. Ashford, T. Suppes, S. Z. Chao, J. Taylor, A. C. Rosen, T. C. Durazzo, L. C. Lazzeroni, J. Yesavage, Repetitive Transcranial Magnetic Stimulation as a Treatment for Veterans with Cognitive Impairment and Multiple Comorbidities. J Alzheimers Dis 85, 1593–1600 (2022).

61. M. Cotelli, M. Calabria, R. Manenti, S. Rosini, O. Zanetti, S. F. Cappa, C. Miniussi, Improved language performance in Alzheimer disease following brain stimulation. J Neurol Neurosurg Psychiatry 82, 794–797 (2011).

62. W. Jiang, Z. Wu, L. Wen, L. Sun, M. Zhou, X. Jiang, Y. Gui, The Efficacy of High-or Low-Frequency Transcranial Magnetic Stimulation in Alzheimer’s Disease Patients with Behavioral and Psychological Symptoms of Dementia. Adv Ther 39, 286–295 (2022).

63. M. Kumar, C. T. Ellis, Q. Lu, H. Zhang, M. Capotă, T. L. Willke, P. J. Ramadge, N. B. Turk-Browne, K. A. Norman, BrainIAK tutorials: User-friendly learning materials for advanced fMRI analysis. PLoS Comput Biol 16, e1007549 (2020).

64. H. Li, J. Ma, J. Zhang, W.-Y. Shi, H.-N. Mei, Y. Xing, Repetitive Transcranial Magnetic Stimulation (rTMS) Modulates Thyroid Hormones Level and Cognition in the Recovery Stage of Stroke Patients with Cognitive Dysfunction. Med Sci Monit 27, e931914 (2021).

65. B. J. Lithgow, Z. Dastgheib, Z. Moussavi, Baseline Prediction of rTMS efficacy in Alzheimer patients. Psychiatry Res 308, 114348 (2022).

66. H. Lu, S. S. M. Chan, S. Ma, C. Lin, V. C. T. Mok, L. Shi, D. Wang, A. D.-P. Mak, L. C. W. Lam, Clinical and radiomic features for predicting the treatment response of repetitive transcranial magnetic stimulation in major neurocognitive disorder: Results from a randomized controlled trial. Hum Brain Mapp 43, 5579–5592 (2022).

67. P. R. Padala, E. M. Boozer, S. Y. Lensing, C. M. Parkes, C. R. Hunter, R. A. Dennis, R. Caceda, K. P. Padala, Neuromodulation for Apathy in Alzheimer’s Disease: A Double-Blind, Randomized, Sham-Controlled Pilot Study. Journal of Alzheimer’s Disease 77, 1483–1493 (2020).

68. Y. Qin, F. Zhang, M. Zhang, W. Zhu, Effects of repetitive transcranial magnetic stimulation combined with cognitive training on resting-state brain activity in Alzheimer’s disease. Neuroradiol J 35, 566–572 (2022).

69. G. Rutherford, B. Lithgow, Z. Moussavi, Short and Long-term Effects of rTMS Treatment on Alzheimer’s Disease at Different Stages: A Pilot Study. J Exp Neurosci 9, 43–51 (2015).

70. Y. Saitoh, K. Hosomi, T. Mano, Y. Takeya, S. Tagami, N. Mori, A. Matsugi, Y. Jono, H. Harada, T. Yamada, A. Miyake, Randomized, sham-controlled, clinical trial of repetitive transcranial magnetic stimulation for patients with Alzheimer’s dementia in Japan. Front Aging Neurosci 14, 993306 (2022).

71. Y. Tao, B. Lei, Y. Zhu, X. Fang, L. Liao, D. Chen, C. Gao, Repetitive Transcranial Magnetic Stimulation Decreases Serum Amyloid-β and Increases Ectodomain of p75 Neurotrophin Receptor in Patients with Alzheimer’s Disease. J Integr Neurosci 21, 140 (2022).

72. P. Turriziani, D. Smirni, G. R. Mangano, G. Zappalà, A. Giustiniani, L. Cipolotti, M. Oliveri, Low-Frequency Repetitive Transcranial Magnetic Stimulation of the Right Dorsolateral Prefrontal Cortex Enhances Recognition Memory in Alzheimer’s Disease. J Alzheimers Dis 72, 613–622 (2019).

73. Y. Wu, W. Xu, X. Liu, Q. Xu, L. Tang, S. Wu, Adjunctive treatment with high frequency repetitive transcranial magnetic stimulation for the behavioral and psychological symptoms of patients with Alzheimer’s disease: a randomized, double-blind, sham-controlled study. Shanghai Arch Psychiatry 27, 280–288 (2015).

74. X. Zhang, H. Ren, Z. Pei, C. Lian, X. Su, X. Lan, C. Chen, Y. Lei, B. Li, Y. Guo, Dual-targeted repetitive transcranial magnetic stimulation modulates brain functional network connectivity to improve cognition in mild cognitive impairment patients. Front Physiol 13, 1066290 (2022).

75. X. Zhou, Y. Wang, S. Lv, Y. Li, S. Jia, X. Niu, D. Peng, Transcranial magnetic stimulation for sleep disorders in Alzheimer’s disease: A double-blind, randomized, and sham-controlled pilot study. Neurosci Lett 766, 136337 (2022).

76. S. L. Thompson-Schill, M. D’Esposito, G. K. Aguirre, M. J. Farah, Role of left inferior prefrontal cortex in retrieval of semantic knowledge: A reevaluation. Proceedings of the National Academy of Sciences 94, 14792–14797 (1997).

77. A. D. Wagner, E.J. Paré-Blagoev, J. Clark, R. A. Poldrack, Recovering Meaning: Left Prefrontal Cortex Guides Controlled Semantic Retrieval. Neuron 31, 329–338 (2001).

78. A. H. Kizilbash, R. D. Vanderploeg, G. Curtiss, The effects of depression and anxiety on memory performance. Archives of Clinical Neuropsychology 17, 57–67 (2002).

79. F. Manes, R. Jorge, M. Morcuende, T. Yamada, S. Paradiso, R. G. Robinson, A controlled study of repetitive transcranial magnetic stimulation as a treatment of depression in the elderly. Int Psychogeriatr 13, 225–231 (2001).

80. P. Sabesan, S. Lankappa, N. Khalifa, V. Krishnan, R. Gandhi, L. Palaniyappan, Transcranial magnetic stimulation for geriatric depression: Promises and pitfalls. World J Psychiatry 5, 170–181 (2015).

81. M. Corbetta, L. Ramsey, A. Callejas, A. Baldassarre, C. D. Hacker, J. S. Siegel, S. V. Astafiev, J. Rengachary, K. Zinn, C. E. Lang, L. T. Connor, R. Fucetola, M. Strube, A. R. Carter, G. L. Shulman, Common Behavioral Clusters and Subcortical Anatomy in Stroke. Neuron 85, 927–941 (2015).

82. B. T. Thomas Yeo, F. M. Krienen, J. Sepulcre, M. R. Sabuncu, D. Lashkari, M. Hollinshead, J. L. Roffman, J. W. Smoller, L. Zöllei, J. R. Polimeni, B. Fischl, H. Liu, R. L. Buckner, The organization of the human cerebral cortex estimated by intrinsic functional connectivity. Journal of Neurophysiology 106, 1125–1165 (2011).

83. A. Cohen, L. Soussand, P. McManus, M. Fox, GSP1000 Preprocessed Connectome (2021), doi:10.7910/DVN/ILXIKS.

84. B. Milner, Visual recognition and recall after right temporal-lobe excision in man. Neuropsychologia 6, 191–209 (1968).

85. D. Kimura, Perception of Unfamiliar Stimuli After Damage. Archives of Neurology 8, 264–271.

86. R. H. B. Benedict, D. Schretlen, L. Groninger, M. Dobraski, B. Shpritz, Revision of the Brief Visuospatial Memory Test: Studies of normal performance, reliability, and validity. Psychological Assessment 8, 145–153 (1996).

87. E. Grober, R. B. Lipton, R. A. Sperling, K. V. Papp, K. A. Johnson, D. M. Rentz, A. E. Veroff, P. S. Aisen, A. Ezzati, Associations of Stages of Objective Memory Impairment With Amyloid PET and Structural MRI. Neurology 98, e1327–e1336 (2022).

88. A. Thielscher, A. Antunes, G. B. Saturnino, in 2015 37th Annual International Conference of the IEEE Engineering in Medicine and Biology Society (EMBC), (2015), pp. 222–225.

89. P. H. Westfall, S. S. Young, Resampling-Based Multiple Testing: Examples and Methods for p-Value Adjustment (John Wiley & Sons, 1993).

